# When will the cancer start? Elucidating the correlations between cancer initiation times and lifetime cancer risks

**DOI:** 10.1101/640284

**Authors:** Hamid Teimouri, Maria Kochugaeva, Anatoly B. Kolomeisky

**Affiliations:** Department of Chemistry, Rice University, Houston, Texas, United States; Center for Theoretical Biological Physics, Rice University, Houston, Texas, United States; Systems Biology Institute, Yale University, West Haven, CT, United States; Department of Chemical and Biomolecular Engineering, Rice University, Houston, Texas, United States; Department of Physics and Astronomy, Rice University, Houston, TX, United States

## Abstract

Cancer is a genetic disease that results from accumulation of unfavorable mutations. As soon as genetic and epigenetic modifications associated with these mutations become strong enough, the uncontrolled tumor cell growth is initiated, eventually spreading through healthy tissues. Clarifying the dynamics of cancer initiation is thus critically important for understanding the molecular mechanisms of tumorigenesis. Here we present a new theoretical method to evaluate the dynamic processes associated with the cancer initiation. It is based on a discrete-state stochastic description of the formation of tumors as a fixation of unfavorable mutations in tissues. Using a first-passage analysis the probabilities for the cancer to appear and the times before it happens, which are viewed as fixation probabilities and fixation times, respectively, are explicitly calculated. It is predicted that the slowest cancer initiation dynamics is observed for neutral mutations, while it is fast for both advantageous and, surprisingly, disadvantageous mutations. The method is applied for estimating the cancer initiation times from experimentally available lifetime cancer risks for different types of cancer. It is found that the higher probability of the cancer to occur does not necessary lead to the fast times of starting the cancer. Our theoretical analysis helps to clarify microscopic aspects of cancer initiation processes.

## 1 Introduction

It is well known that tumor cells are characterized by abnormal cell division rates, which is a result of mutations in cancer-susceptible genes (known as oncogenes) [5,8,10,22]. Specifically, these mutations affect the regulation of cell proliferation and differentiation via activation of oncogenes or inactivation of tumor suppressor genes (TSGs) [5,8,19,22]. Mutations are taking place randomly, and after several cellular replications some of them might occasionally lead to significant genetic and epigenetic alterations such that the normal cells behavior changes to the uncontrolled proliferation, eventually starting a cancer [1,7,8]. After these cancer initiation events, rapid changes are taking place with a newly formed tumor being able to escape cellular control mechanisms, and the cancer progresses into more invasive forms [5, 7, 8, 17, 18]. But this happens only after the initial stage of cancer succeeds, and thus it is critically important to understand the dynamics of cancer initiation [7].

Human tissues and organs are composed of heterogeneous mixtures of cells: not all cells are equal in their potential to proliferate. An important role in tissue maintenance and repair is played by a population of so-called stem cells [31]. These cells are characterized by their ability to self-renew and make more stem cells or ability to produce differentiated progenitor cells [2]. Epithelial tissues are also known for subdivision into compartments where homeostatic mechanism, a balance between self-renewal and differentiation, maintains the constant cell number. Cancer appears in such compartments, breaking the homeostatic tissue equilibrium. However, having only a single mutated cell in the compartment does not lead to cancer. The cancer initiation event generally is associated with a fixation of one or several mutations, i.e, when all cells in the compartment become mutated, or when a significant fraction of them is mutated, producing noticeable genetic and epigenetic changes [7,17].

One of the most important quantities that determines if the person gets a cancer is a cancer lifetime risk. It refers to a probability of being diagnosed with or dying from cancer during the person’s lifespan. Lifetime risks strongly depend on the type of cancer. For example, a person’s risk of getting a lung cancer is more than 11 times that of developing of a brain cancer, and 8 times greater than that of a stomach cancer [6, 29]. Various studies have attributed the differences in cancer rates to environmental risk factors, such as smoking, bad dietary habits or exposure to UV light, as well as to heritable mutations. However, the environmental factors and the hereditary factors cannot fully explain the substantial differences in the cancer rates across tissues. Moreover, the total numbers of cells that make up these tissues also cannot explain varying cancer risks. Recent statistical analysis of 31 cancers by Tomesetti and Vogelstein suggested that there is strong correlation between random mutations acquired during stem cell divisions and lifetime cancer risk [28, 29].

It is widely assumed that the cancer initiation time is inversely proportional to the lifetime cancer risk, i.e, the higher the lifetime risk, the shorter is the time before the cancer starts. However, this issue has not been methodically investigated. There are certain types of cancers with low lifetime risks that occur at early ages, while there are other types with high lifetime risks that happen at older ages. Therefore it is crucially important to estimate the initiation times for different cancer types. In recent years, several mathematical models have been developed for analysis of cancer initiation and progression dynamics [1, 3, 5, 7, 9, 14, 19, 22, 27]. However, some important microscopic aspects of the evolutionary processes leading to cancer remains unexplained. For example, the state of the system when the mutated cells take over the whole tissue compartment, which is known as a fixation, is frequently associated with the formation of the tumor [22]. While the probability to reach the fixation, called a fixation probability, has been explicitly evaluated [22], the time to reach the fixation (fixation time) have been estimated only approximately using numerical and computer simulations methods [11, 24].

Here we develop a new theoretical framework of explicitly evaluating the cancer initiation dynamics. The mutation fixation in the tissue compartment is assumed as the point of cancer initiation. Applying a discrete-state stochastic approach with a first-passage analysis, the fixation probabilities and fixation times are calculated exactly. Utilizing our theoretical predictions we extract relevant parameters from experimental data on lifetime risks for different types of cancer, which are used then to estimate the specific cancer initiation times. Our theoretical analysis suggests that there is no correlations between the probability and mean time of getting cancer, suggesting that *both* properties should be utilized in evaluation of cancer risks.

## 2 Methods

### 2.1 Theoretical model

Let us consider a tissue compartment that has at the beginning *N* normal stem cells as shown in Fig. 1 (Top). At some specific time, which we set as a time zero, one stem cell undergoes a mutation with a probability *μ*. Here we consider only driver mutations, i.e. those that promote the cancer development [18, 25]. Both normal and mutated stem cells in the tissue can replicate, but with different rates. To reflect the effect of somatic mutations, the mutated cells are characterized by a fitness parameter *r*, which is defined as the ratio of the division rate of the mutated cells to the division rate of the normal cells. If *r* > 1 the mutation is advantageous; if *r* < 1 the mutation is disadvantageous; and *r* =1 describes a neutral mutation. It is expected that most mutations in oncogenes that lead to cancer are advantageous [22]. The important characteristic of normal tissues is the homeostatic equilibrium, i.e., the total number of cells in the compartment remains constant. To incorporate this property into our theoretical model we assume that the system follows a birth-death process known as a Moran process [20–22]. This means that after division of any randomly chosen cell the number of cells in the compartment increases by one, and then one of the randomly chosen cells should be instantly removed to keep the number of cell constant and equal to *N*: see Fig. 1 (Top). It is also assumed that there is no other mutations in the tissue compartment. This is a reasonable assumption because cell division rates are much faster than the mutation rates for driver mutations [22, 25]. Because there are two types of cells in the tissue compartment. each of the state of the system can be labeled as *n*, where *n* is the number of mutated cells and *N* — *n* is the number of normal cells. Then all transformations in the system can be viewed as random transitions between corresponding discrete states as presented in Fig. 1 (Bottom). After the mutation happens, the system can get rid of this mutation - this corresponds to going from the state 1 to the state 0. But the number of mutation can also increase and eventually the system might reach the state *N*, which corresponds to mutation fixation (see Fig. 1). We identify the fixation state as a starting point of the cancer because there are no normal stem cells left in the compartment [5,22]. Thus, the cancer initiation dynamics in our model is viewed as a process of transitioning from the state 1 to the state *N*. This is a stochastic process which is governed by various transition rates. Following the description of the Moran processes [22], and considering the two-stage process for replication via cell division and immediate cell removal to fix the total number of cells (see the details of derivation in the SI), the forward transition rate from the state *n* to the state *n* + 1 is given by *ra*_*n*_ where

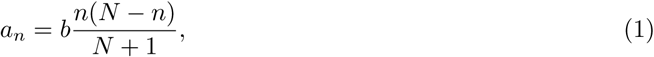

and *b* is a division rate of the normal stem cell. The factor *r* comes from the fact that this transition is taking place due to replication of the mutated cell and the corresponding instantaneous removal (to keep the homeostatic equilibrium) of the normal stem cell. The backward transition (from the state *n* to the state *n* — 1) is equal just to *a*_*n*_ because it describes the replication of the normal cell and the removal of the mutated cell. In our theoretical framework, the cancer starts when the system reaches the state *N* for the first time starting initially in the state 1. This suggests that the cancer initiation dynamics can be conveniently described as a first-passage process [13,26]. One can define a first-passage probability density function *F*_*n*_ (*t*) that describes the probability to reach the state *N* for the first time at time *t* if at *t* = 0 the system started in the state *n*. The temporal evolution of these functions can be described by a set of so-called backward master equations [13, 26]

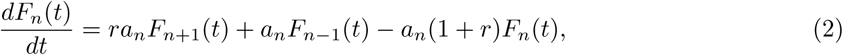

for 1 < *n* < *N*; and

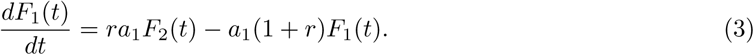

**Figure 1.**
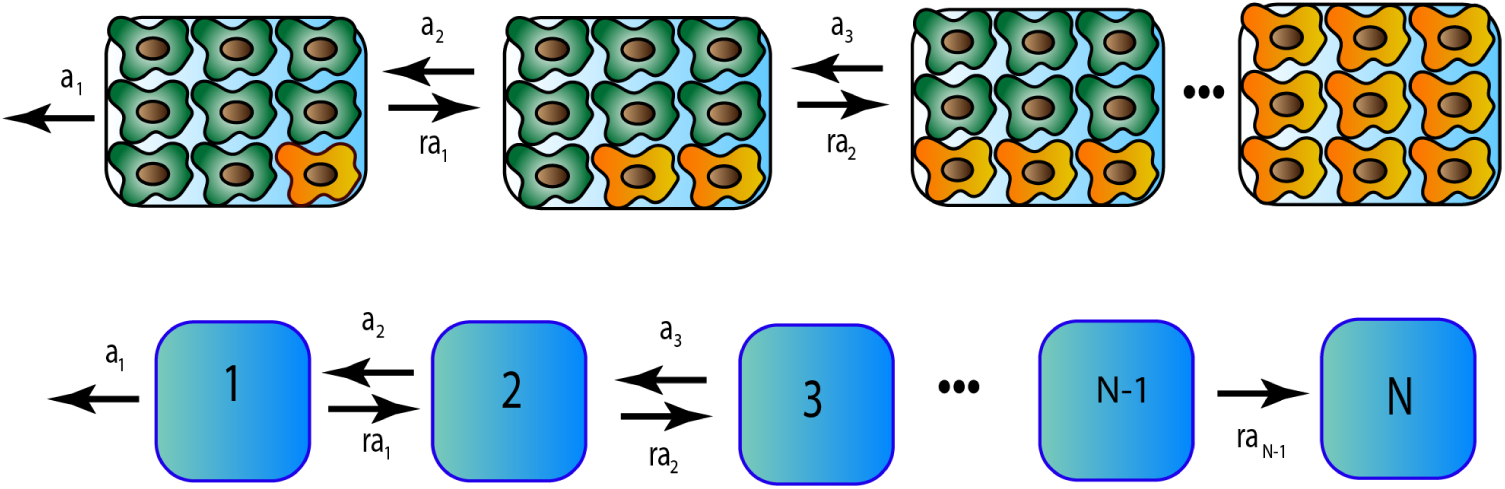
Top: A schematic view of a single mutation fixation process in the tissue compartment. Normal stem cells are green, while mutated cells are yellow. Bottom: Corresponding discrete-state stochastic model.

In addition, we have the boundary condition *F*_*N*_(*t*) = *δ(t)*, the physical meaning of which is that the fixation process is immediately accomplished if the system starts from the state *N*.

First-passage probability functions contain a comprehensive dynamic description of the fixation process. In this work, we are interested in fixation probabilities 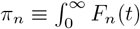 and fixation times 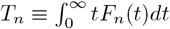, which are analytically calculated in the Appendix. For example, for the fixation probability from the state *n* we obtain

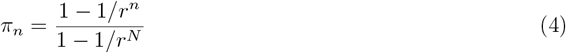

which is a well known result [22]. For *r* → 1 we get *π*_*n*_ = *n/N*. In Fig. 2a, the fixation probability *π*_1_ is presented for different values of the parameters *r* and *N*. One can see that for large values of *N* the fixation probability depends only on the fitness parameter *r*.

**Figure 2.**
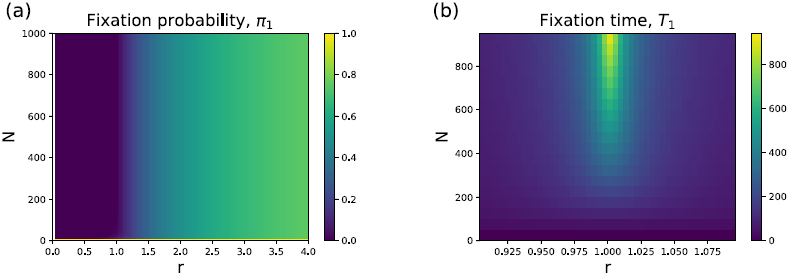
Heat maps for (a) fixation probability *π*_1_ and (b) fixation time *τ*_1_ (normalized with respect to the normal stem cell replication time, i.e, *b* = 1) as a function parameters *r* and *N*.

A critically important feature of the cancer initiation process is how long does it take to reach the cancer starting point. In our language, it corresponds to the fixation time for the mutation that activated the oncogene [3, 24]. More specifically, it is given by *T*_1_, which is as a conditional mean first-passage time to reach the fixation state *(n* = *N*) from the state with initially *n* = 1 mutated cells before the mutation can be eliminated from the system *(n* = 0). Our explicit calculations (presented in the SI) provide the following expression,

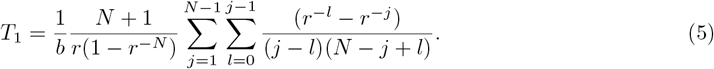

For *r* →1 and *N* →∞ we obtain:

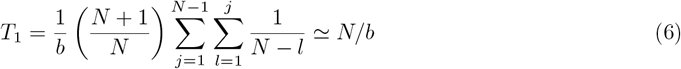

It can be also shown (see SI for details) that for large *N* the expression for the fixation time can be simplified into

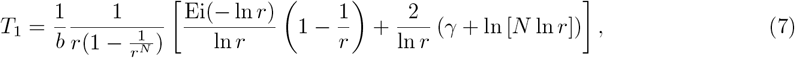

where *Ei(x)* is the exponential integral defined as 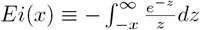, and *γ* is the Euler-Mascheroni constant. The results for fixation times as functions of *N* and *r* are presented in Fig. 2b. The slowest cancer initiation dynamics is expected for the neutral mutations (*r* = 1). This can be easily understood because in this case the system performs the unbiased random walk between the discrete states (see Fig. 1 Bottom), redundantly visiting the same states many times. As expected, for the advantageous mutations (*r* > 1) the cancer initiation times are lower since the dynamics is biased in the direction of increasing the number of mutated cells in the tissue compartment (Fig. 1). This will drive the system faster in the direction of the fixation. Surprisingly, the fixation times are also fast for the disadvantageous mutations (*r* < 1). This unexpected result can be explained using the following arguments. The system is biased in the direction of decreasing the number of mutations and this leads to very low fixation probabilities (see Fig. 2a). However, for those rare events when the system goes to the fixation they must happen very fast in order not to be influenced by the bias in the opposite direction.

### 2.2 Estimation of fitness parameters for different types of cancer

To calculate explicitly the initiation times for different types of cancer, we need to estimate the fitness parameter *r* and the number of stem cells *N* in the specific tissues. The latter has been well evaluated in Ref. [29] (see Fig. 3). However, the estimation of the fitness parameter *r* is much more difficult, and it requires several approximations.

**Figure 3.**
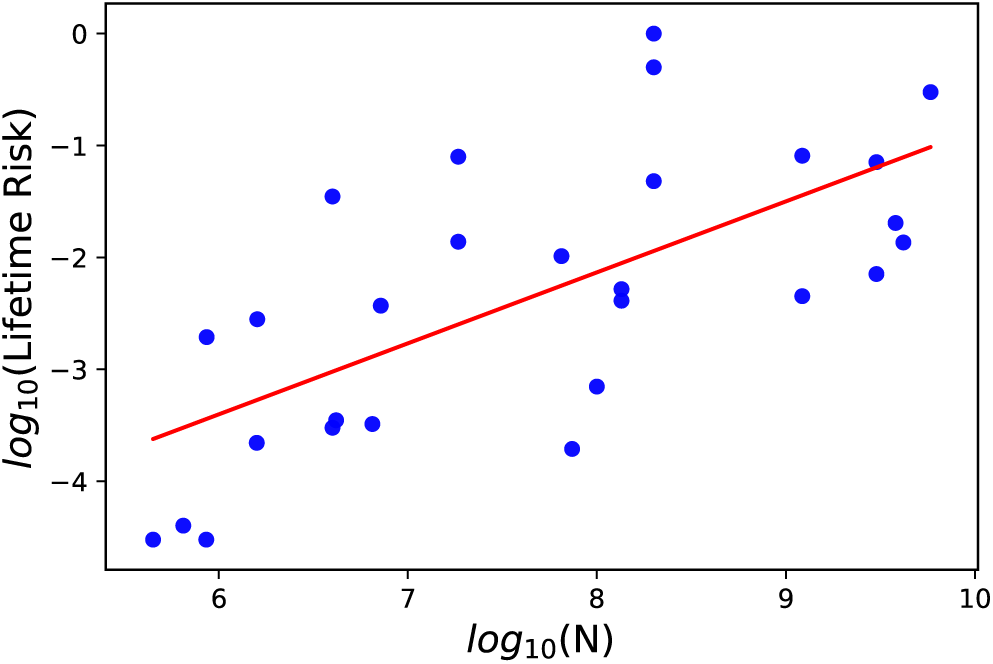
The cancer lifetime risk (*R*_*ltr*_) as a function of the initial number of stem cells (*N*). Correlation analysis yields a Spearman’s correlation coefficient of 0.72. Data are taken from [29].

In our analysis, we follow a simple mathematical approach proposed in Ref. [23]. According to this method, the cancer initiation probability, i.e., the probability that a mutation is fixed in the compartment of *N* cells during a human lifetime, is given by:

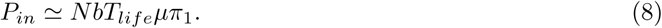

where *T*_*life*_ is the typical human lifetime (we assume here *T*_*life*_ = 80 years), *μ* is the probability of mutation (activation of the oncogene) multiplied by the number of possible oncogenes, which varies for the different tissues [16]. This result can be physically explained using the following arguments. The system can move in the direction of the fixation state, which is associated with the start of the cancer, only after cell divisions are taking place. There are *Nb* such divisions per unit time in the tissue with N cells, and over the human lifetime the total number of such divisions will be *Nb*_*life*_. The cancer will not start until, at least, one of the oncogenes is activated, which has the probability *μ*. Finally, *π*_1_ describes the fixation probability that this mutation will not disappear but will occupy the whole tissue compartment.

However, the cancer initiation probability *P*_*in*_ is not exactly the cancer lifetime risk *R*_*ltr*_ that have been evaluated from various clinical data. At the same time, it can be argued that both quantities are related as [23]

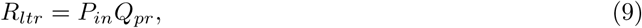

where *Q*_*pr*_ is a probability of cancer progression, i.e., the probability that after the cancer initiation the tumor will grow and the homeostatic equilibrium will be broken. From Eqs. (8) and (9), we obtain

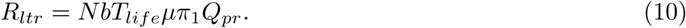

Another factor that helps in estimating the fitness parameter *r* is that, typically, the number of stem cells *N* is very large. This leads to

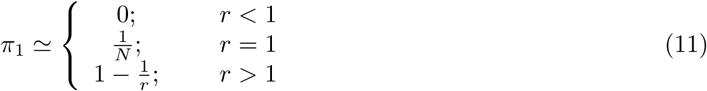

We combine Eqns 10 and 11, and this yields

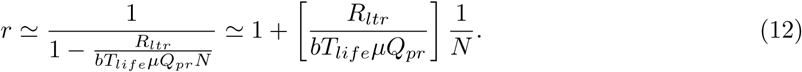

This is an important result because it relates the fitness parameter to the number of stem cells. It is also consistent with ideas presented in Ref. [7], where it was argued that cancer initiation corresponds to gaining a fitness value greater that the threshold value *r\** = 1 + 1/*N*. Finally, Eq. (12) was used to estimate the fitness parameters for several types of cancers, and the results are presented in Table 1.

**Table 1.** Cancer development properties for 28 different cancer types. Data are adapted from [29]. In calculations of fixation times and *t*_*0*_ *μ*= 3 × 10^−8^ and *Q*_*pr*_ = 0.001 were utilized.

| Cancer type | Lifetime risk, $R_{ltr}$ | $N$ | Division rate, $b$ (per year) | $r$ | Fixation time, $T_1$ (years) | $t_0$ (years) |
| --- | --- | --- | --- | --- | --- | --- |
| Acute myeloid leukemia | 0.0041 | $1.35 \times 10^8$ | 12.00 | 1.0011 | 1964.0 | 0.021 |
| Basal cell carcinoma | 0.3000 | $5.82 \times 10^9$ | 7.60 | 1.0028 | 1595.8 | 0.001 |
| Chronic lymphocytic leukemia | 0.0052 | $1.35 \times 10^8$ | 12.00 | 1.0013 | 1577.6 | 0.021 |
| Colorectal adenocarcinoma | 0.0480 | $2.00 \times 10^8$ | 73.00 | 1.0014 | 261.5 | 0.002 |
| Colorectal adenocarcinoma with FAP | 1.0000 | $2.00 \times 10^8$ | 73.00 | 1.0285 | 15.0 | 0.002 |
| Colorectal adenocarcinoma with lynch syndrome | 0.5000 | $2.00 \times 10^8$ | 73.00 | 1.0143 | 29.2 | 0.002 |
| Duodenum adenocarcinoma | 0.0003 | $4.00 \times 10^6$ | 24.00 | 1.0013 | 583.6 | 0.35 |
| Duodenum adenocarcinoma with FAP | 0.0350 | $4.00 \times 10^6$ | 24.00 | 1.1519 | 6.6 | 0.35 |
| Esophageal squamous cell carcinoma | 0.0019 | $8.64 \times 10^5$ | 17.40 | 1.0537 | 22.9 | 2.2 |
| Gallbladder non papillary adenocarcinoma | 0.0028 | $1.60 \times 10^6$ | 0.58 | 2.2486 | 18.3 | 35.7 |
| Head & neck squamous cell carcinoma | 0.0138 | $1.85 \times 10^7$ | 21.50 | 1.0145 | 82.8 | 0.08 |
| Head & neck squamous cell carcinoma with HPV-16 | 0.0794 | $1.85 \times 10^7$ | 21.50 | 1.0831 | 15.2 | 0.08 |
| Hepatocellular carcinoma | 0.0071 | $3.01 \times 10^9$ | 0.91 | 1.0011 | 31640.3 | 0.012 |
| Hepatocellular carcinoma with HCV | 0.0710 | $3.01 \times 10^9$ | 0.91 | 1.0108 | 3593.6 | 0.012 |
| Lung adenocarcinoma (nonsmokers) | 0.0045 | $1.22 \times 10^9$ | 0.07 | 1.0220 | 22468.1 | 0.39 |
| Lung adenocarcinoma (smokers) | 0.0810 | $1.22 \times 10^9$ | 0.07 | 1.3952 | 1058.7 | 0.39 |
| Melanoma | 0.0203 | $3.80 \times 10^9$ | 2.48 | 1.0009 | 14019.2 | 0.004 |
| Osteosarcoma | 0.0004 | $4.18 \times 10^6$ | 0.07 | 1.5207 | 566.6 | 119.02 |
| Osteosarcoma of the arms | 0.00004 | $6.50 \times 10^5$ | 0.07 | 1.3827 | 725.5 | 765.4 |
| Osteosarcoma of the head | 0.00003 | $8.60 \times 10^5$ | 0.07 | 1.2169 | 1423.3 | 578.5 |
| Osteosarcoma of the legs | 0.00022 | $1.59 \times 10^6$ | 0.07 | 1.8605 | 272.0 | 312.9 |
| Osteosarcoma of the pelvis | 0.00003 | $4.50 \times 10^5$ | 0.07 | 1.4146 | 639.9 | 1105.6 |
| Pancreatic ductal adenocarcinoma | 0.0136 | $4.18 \times 10^9$ | 1.00 | 1.0014 | 23772.1 | 0.008 |
| Pancreatic endocrine carcinoma | 0.0002 | $7.40 \times 10^7$ | 1.00 | 1.0011 | 21716.2 | 0.45 |
| Small intestine adenocarcinoma | 0.0007 | $1.00 \times 10^8$ | 36.00 | 1.0001 | 6566.2 | 0.009 |
| Testicular germ cell cancer | 0.0037 | $7.20 \times 10^6$ | 5.80 | 1.0369 | 117.3 | 0.798 |
| Thyroid papillary/follicular carcinoma | 0.0103 | $6.50 \times 10^7$ | 0.09 | 1.7560 | 314.5 | 5.9 |
| Thyroid medullary carcinoma | 0.0003 | $6.50 \times 10^6$ | 0.09 | 1.2387 | 1143.0 | 58.95 |

### 2.3 Estimation of the fixation times and times before the first mutation appears

After determining the fitness parameter *r*, we can now estimate the cancer initiation times, which in our theoretical framework are the same as the fixation times. Because the number of stem cells is very large, it can be shown from Eqs. (7) and (12) that

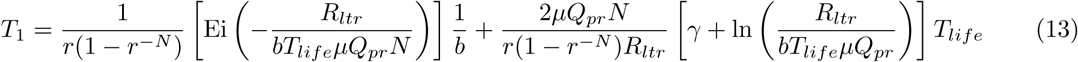

In calculating the fixation times, cancer lifetime risks *R*_*ltr*_, number of stem cells *N* and cell division rates *b* were taken from the data assembled in Ref. [29]. The probability for activating a single oncogene was estimated to be ≃ 3 × 10^−9^ and multiplying it by the average number of oncogenes ~ 10 we obtain *μ* ≃ 3 × 10^−8^ [15, 16, 22]. Much less information is known about the probability of cancer progression (*Q*_*pr*_). It has been argued theoretically and supported by some medical data that not all lesions progress to full cancers [4, 18, 30], and we estimate *Q*_*pr*_ ≃ 0.001. Using our theoretical framework, we can also estimate the time before the first mutation appears, *t*_0_. It can be shown that it is given by

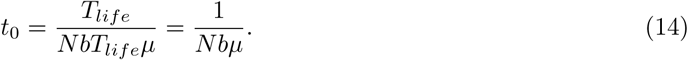

This formula can be understood by noting that *NbT*_*life*_*μ* gives the total number of driver mutations during the lifetime, and dividing the lifetime by this number gives the average time between mutations.

The results of our calculations for the fixation times and for the times before the first mutation appears are presented in Table 1. Our theoretical method suggests that the fixation times vary strongly for different types of cancer. Slightly smaller variations are predicted for *t*_0_. However, one should be cautioned to take these numbers literally because they are sensitive to absolute values of *μ* and *Q*_*pr*_, and we used the same values of *μ* and *Q*_*pr*_ for all cancers just to illustrate our method, which is also not realistic. However, we believe that these calculations provide a reasonable description of trends in the cancer initiation dynamics.

### 2.4 Correlation between cancer lifetime risks and cancer initiation times

The cancer lifetime risks are widely utilized for predicting the chances of getting the cancer. It is also frequently implicitly assumed that the higher the risk, the faster cancer will develop. However, the relations between cancer lifetime risks and cancer initiation times have not been thoroughly investigated. Our theoretical method allows us to measure the correlations between these quantities because both properties can be explicitly evaluated.

Fig. 4 shows the fixation times, estimated using our method, and experimental data on cancer lifetime risks for 28 different types of cancer from Ref. [28]. Statistical analysis of this graph gives a Spearman’s correlation coefficient −0.2 between the cancer lifetime risks and the fixation times, magnitude of which is significantly smaller than the value −1 expected for the perfect correlation. To test the validity of the null hypothesis that there is no correlation, we also performed a p-value analysis of these data. A large p-value of *p* = 0.31 supports the null hypothesis. Thus, we predict that there is no correlations between cancer lifetime risks and cancer initiation times. This is a very important result because it argues that cancer lifetime risks alone cannot be utilized to evaluate the danger of getting cancer. Cancer initiation times should also be utilized, and we provide the quantitative framework how to estimate them.

**Figure 4.**
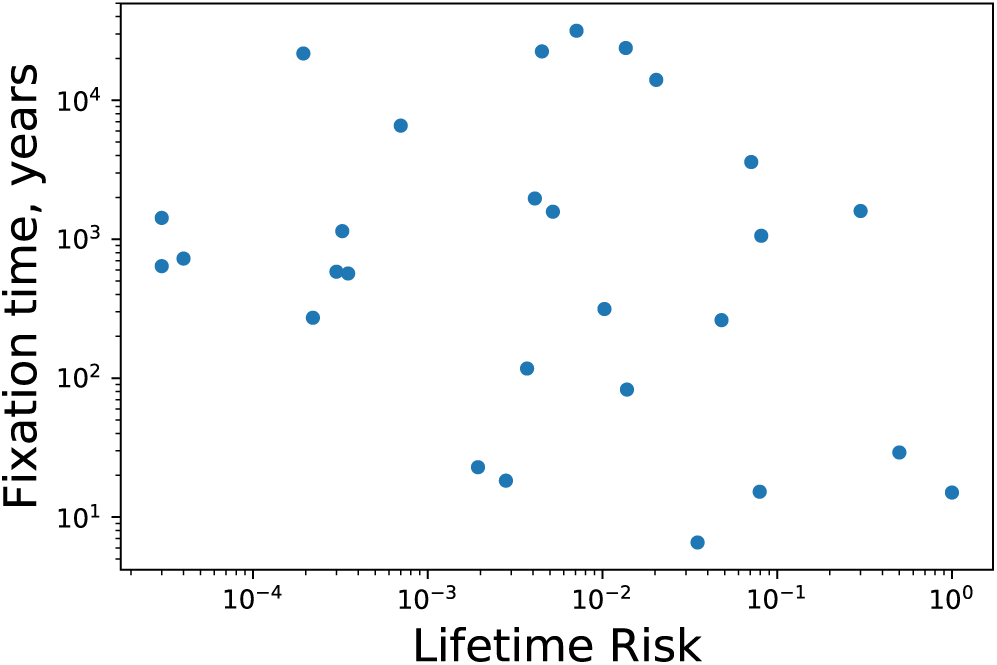
Fixation time vs lifetime risk for different types of cancer.

## 3 Discussion

We developed a simple mathematical approach to evaluate the cancer initiation dynamics. The appearance of tumor is associated with fixation of some random mutation in the originally healthy tissue with fixed number of stem cells. The initial stage of cancer development is viewed as a stochastic process of transitions between discrete states with different numbers of mutated cells, and the first-passage analysis is utilized for calculating exactly the fixation probabilities and the fixation times.

It is shown that the cancer initiation dynamics depends strongly on the fitness parameter r that describes how faster the mutated cell divides in comparison with the normal cell. The effect of the number of cells N in the tissue is much weaker. It is found that for large fitness parameters the probability of fixation, as expected. However, the dependence of fixation times is non-monotonic with the maximum for neutral mutations (*r* = 1). What is surprising that even for disadvantageous mutations (*r* < 1) the fixation might start quite quickly. We are able to explain these observations by utilizing arguments that view the fixation process as a random walk on the sequence of states with different degrees of mutations. Neutral mutations correspond to the unbiased random walk, which is slow because many states are repeatedly visited during the process. For disadvantageous mutations, which can be viewed as a biased random walk in the direction opposite to the fixation, the probability of fixation is small. Then only those events lead to the fixation that are fast enough so that the bias does not have time to act.

We applied our theoretical approach for evaluating explicitly the initiation times for different types of cancer. This is done by connecting theoretically calculated fixation probabilities with available clinical data on cancer lifetime risks, from which the fitness parameters are estimated. This allows us to calculate exactly the fixation times that are associated with the starting point of the cancer. The initiation times are determined for 28 different types of cancer. We performed then the analysis of correlations between cancer lifetime risks and the cancer starting times. In contrast to expectations, it is found that there is no correlations between these properties of cancer initiation dynamics, assuming that our theoretical method correctly predicts the starting times for cancer. This has an important consequence suggesting that both dynamic features, lifetime risks and initiation times, are required to comprehensively evaluate the risks of getting cancer.

While our theoretical method cannot provide molecular details to explain the observation on the lack of correlations, we can give the following microscopic arguments using the analogy with thermodynamics and kinetics of chemical processes. It is known that although thermodynamics gives the probability for the chemical reaction to occur, only kinetics determines if the reaction actually takes place on experimentally observable time scales. Thermodynamic probability is proportional to an equilibrium constant for the process, which is the ratio of forward over the backward reaction rates. At the same time, kinetics is determined by the slowest transition rates. In our theoretical language, the fixation probability depends on the product of ratios of forward to backward transition rates between all sequential discrete states (see Fig. 1), which is equal to the fitness parameter r. However, the fixation times depend more on the slowest forward transition rates, which are *ra*_1_ (from the sate 1 to the state 2) and *ra*_*N*−1_ (from the state *N* − 1 to the state *N*). The transition from the state 1 is slow because there is only 1 mutated cell in the tissue. The transition from the state N − 1 to the fixation is slow because only one normal cell left and the probability that it will be picked out for removal is very low. Thus, the slow speed of initial and final transitions during the mutation fixation process might be the reason for general lack of correlations between the fixation probabilities and fixation times.

It is also important to critically evaluate our theoretical method since it involves several approximations and assumptions. We assume that after a single mutation appears in the tissue no other mutations can occur in the system. This is probably a reasonable assumption because the probability of activating the oncogenes *μ* is typically very low and normal cell division rates are typically fast [16, 25, 28]. But if the second mutation appears before the fixation of the first one, it is expected that the overall fixation time should be lower. Multiple studies also suggest that more than one mutation in tumor-suppressing genes (“hits”) is required for cancer to start [12, 19, 22]. Our theoretical approach can be extended to analyze these “two-hit” models. It is expected that while the fixation times will be longer in this case, other qualitative trends should remain the same as discussed here. One could also notice that the explicit values of the fixation times depend on the probability of mutation during the replication *μ* and the probability of cancer progression *Q*_*pr*_. Because both of these parameters are not well determined and depend on the type of the cancer, we varied them by several orders of magnitude (see details in the Appendix). It is found that the magnitude of initiation times are quite sensitive to variations in *μ* and *Q*_*pr*_. In addition, it can be argued that the cancer might start when a large fraction of cells in the tissue, but not all of them, is mutated. Furthermore, the fitness parameter might increase with number of cell replications, and this is also not accounted in our model. Both these effects will shorten the cancer initiation times. Our model also does not take into account spatial effects [14]. But we notice that our theoretical framework can be adapted to evaluate the cancer initiation dynamics for some of these more advanced cases.

All these critical comments suggest that it is probably unreasonable to view all the cancer initiation times reported in Table 1 as realistic. This might also explain too large values for some fixation times. However, the trends predicted by our theoretical method should be valid because all data are considered in the similar way. In addition, our theoretical approach gives a convenient, simple and versatile method to evaluate the cancer initiation dynamics, and it is expected that in future better estimates of relevant parameters will make the evaluation of cancer initiation times more precise and reliable. The proposed theoretical method is also useful in designing more quantified approaches in cancer prevention. For example, it can be argued that *t*_0_ + *T*_1_ might be a better time estimate of the age at which the testing of different cancers should start. Furthermore, our theoretical framework is flexible enough to be extended and generalized to include more complex biochemical and biophysical processes that are taking place during the cancer development.

## Acknowledgments

We thank O. Igoshin, K. Vanaja, D. Ostapenko, X. Li and F. Bocci for useful discussions. The work was supported by the Welch Foundation (C-1559), by the NSF (CHE-1664218), and by the Center for Theoretical Biological Physics sponsored by the NSF (PHY-1427654).

## Appendix

In this supporting information we provide details of calculations for the equations in the main text.

### I. Calculating transition rates *a*_*n*_

To compute the transition rate between the states (*n, N* − *n*) (n mutated cells) and (*n* + 1, *N* − *n* − 1) (*n* +1 mutated cells) we consider the cell replication as a two-state process, as shown in Fig. S1. First, the randomly chosen cell is divided and the number of cells in the tissue increases to *N* +1. Then immediately one of the randomly chosen cells is removed to keep the total number of cells equal to *N*. From the state (*n, N* − *n*) (*n* cells are mutated and *N* − *n* are normal) our system goes to an intermediate state (*n* + 1, *N* − *n*). This corresponds to the division by the mutated cell. The rate for this process is equal to *rbn* because there are *n* mutated cells, each of them can divide with the rate *rb*. The reverse transition from the intermediate state (*n* + 1, *N* − *n*) with *N* +1 total number of cells to the state (*n, N* − *n*) with *N* total cell is equal to *A*(*n* +1). Here *A* is the rate of removal of any randomly chosen cell from the system (here we assume that *A* ≫ *b*). From the intermediate state (*n* + 1, *N* − *n*) the system can also go the state (*n* + 1, *N* − *n* − 1 with the rate *A*(*N* − *n*). One can easily evaluate then the effective time to go from the state (*n, N* − *n*) to the state (*n* + 1, *N* − *n* − 1) as [13]

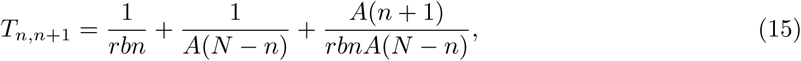

which is equal to the inverse transition rate between these states. From this expression we obtain (recalling that *A* ≫*r*),

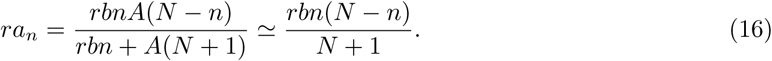

**Appendix I - Figure 1.**
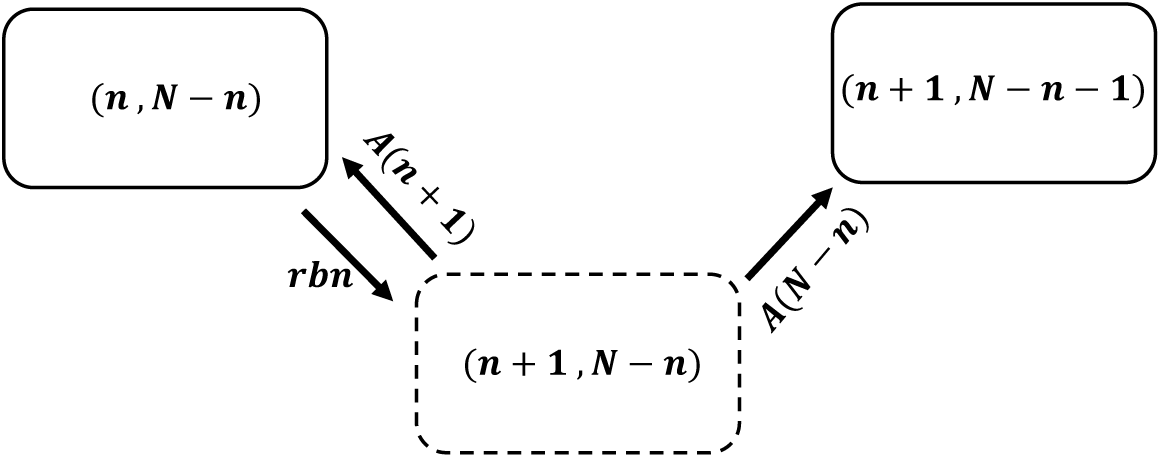
Schematic view for the derivation of eqn 16

### II. Calculation of fixation probability

Let us consider a tissue compartment that has *N* normal cells. At time zero one of them is mutated. Normal cells divide with a speed *b*, while the mutated cell divides with a rate *r* (in units of *b*). Assuming that the compartment always has the same number of cells, let us investigate the dynamics of how the whole compartment can become full of mutant cells. The problem is analogous then to a random walk on the lattice of *N* sites. At *t =* 0 the walk starts at the site 1. The state *n* corresponds to *n* mutated and (*N* − *n*) normal cells. As shown above, the transition rate from the state *n* to *n*+ 1 is equal to *ra*_*n*_, where

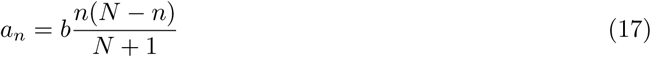

The backward transition (from the state *n* to the state *n* − 1) is equal to *a*_*n*_. The problem of understanding when the whole compartment becomes mutated is analogous then to a first-passage problem of the random walker starting on the site 1 to reach the site *N* for the first time before disappearing to the site 0. One can define the corresponding first-passage probability density functions to start from any site *n* and reach the site *N* at time *t* (if at *t =* 0 the *k* was at the site *n*), *F*_*n*_(*t*). The temporal evolution of these probabilities follows the backward master equations

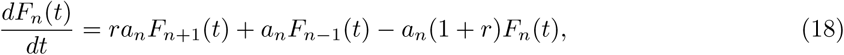

for 1 < *n* < *N;* and

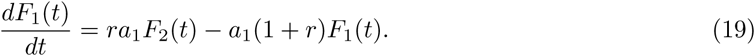

In addition, we have the boundary condition *F*_*N*_(*t*) *= δ*(*t*), which means that the process is immediately accomplished if the walker starts from the site *N*. Let us also do the calculations assuming *b* =1, i.e., all times scales are renormalized with respect to cell replication time.

It is convenient to solve this problem using the Laplace transformation, which changes the backward master equations:

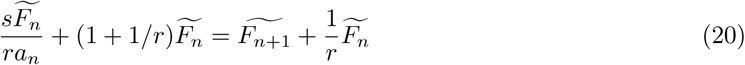

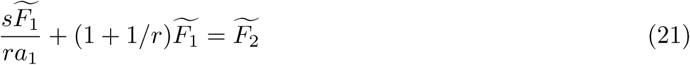

and *F*_*N*_ = 1. Because we are interested only in the fixation probabilities and fixation times, there is no need to obtain full analytical expressions for *F*_*n*_, but it is needed to determine it up to the linear term in s. Thus we can write

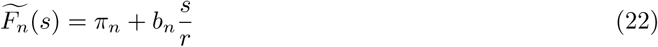

where 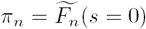, is the fixation probability starting from *n* single mutations, and the unknown parameters *b*_*n*_ are related to the fixation times (viewed as conditional mean first-passage times) as

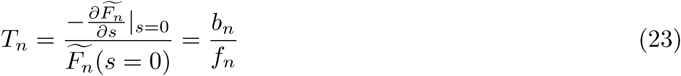

Note that *π*_*N*_ = 1 and *b*_*N*_ = 0. Substituting Eq. 22 into Eqs. 20 and 21 we obtain for the fixation probabilities

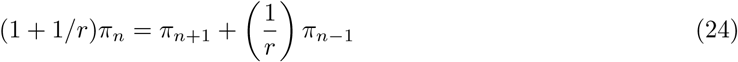

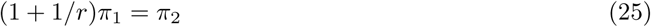

These equations can be easily solved, leading to the following explicit expressions for the fixation probability (a well-known result),

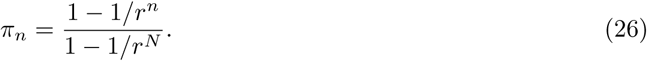

### III. Calculation of fixation time

From Eqs. 20, 21 and 22 the corresponding equations for parameters *b*_*n*_ can be written as,

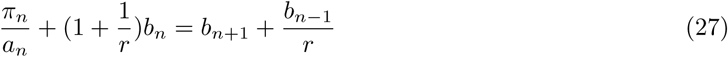

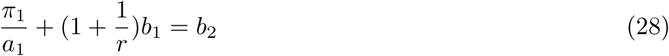

To solve Eqs. 27 and 28 let us write the following anzats

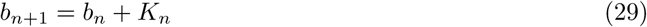

where *K*_*n*_ is another unknown parameter that will be determined. Then the substitution of Eq. 29 into Eqs. 27 and 28 yields

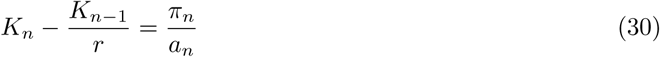

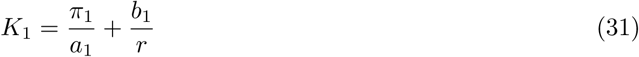

Eq. 30 can be easily solved, producing

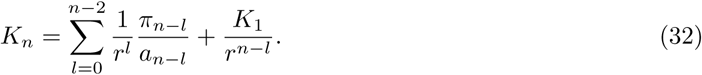

Then from Eq. 29 we can write

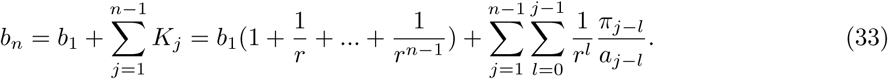

This expression is valid for any 1 ≤ *n* ≤ *N*, then because *b*_*N*_ = 0 we obtain

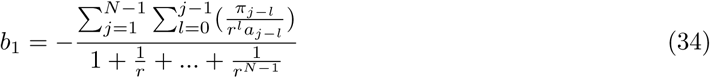

Then the final fixation time (normalized to the cell replication rate *b*) will be equal

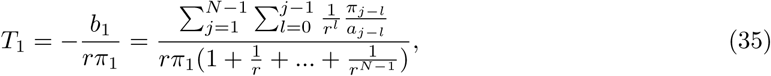

from which after some algebra we obtain

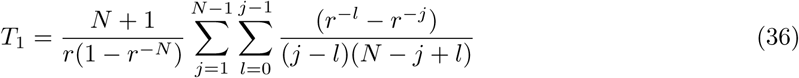

For *r* → 1 and *N* → ∞ we obtain:

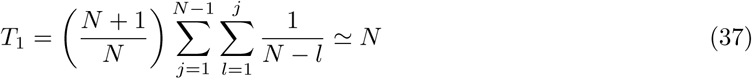

### IV. Explicit expression for fixation times for *N* → ∞

In general it is difficult to perform summation in Eqn. 36. For *N* → ∞, we can convert summation to integration:

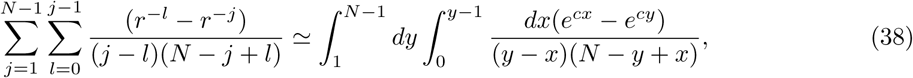

where *c* = − ln *r*. This integral can be written as:

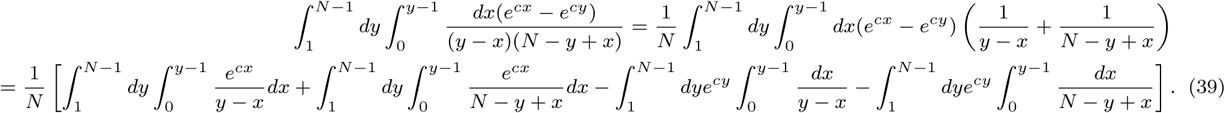

Now we performs integrals term by term:

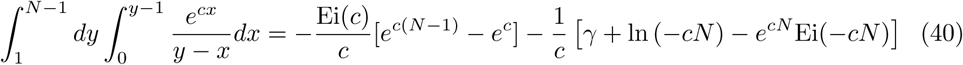

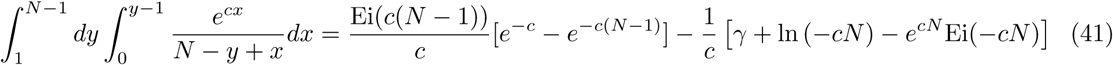

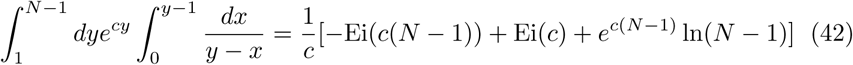

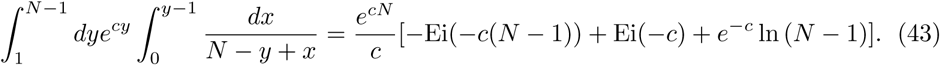

After some algebra, we obtain:

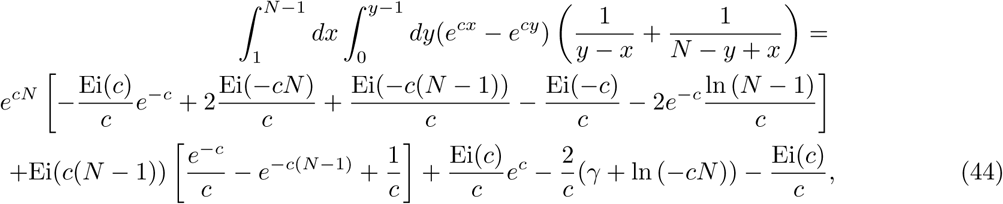

where Ei(x) represents exponential function defined by 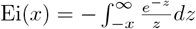, and *γ* is the Euler–Mascheroni constant. Therefore the fixation time is given by

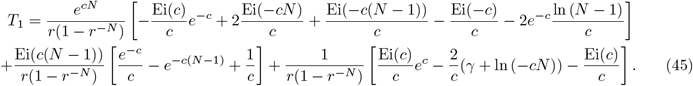

Because *c* < 0 and *N* → ∞, then the first two terms vanish and thus we finally obtain:

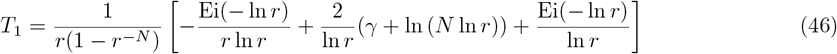

### V. Varying probabilities of cancer progression and oncogene activation

Since our theoretical predictions depend strongly on the probability of cancer progression (*Q*_*pr*_) and the probability of the appearance of mutation (*μ*), which are not well determined in the literature, we varied these parameters as shown in Figure 1. One can see that the calculated fixation times are sensitive to variations in these parameters.

**Appendix V - Figure 1.**
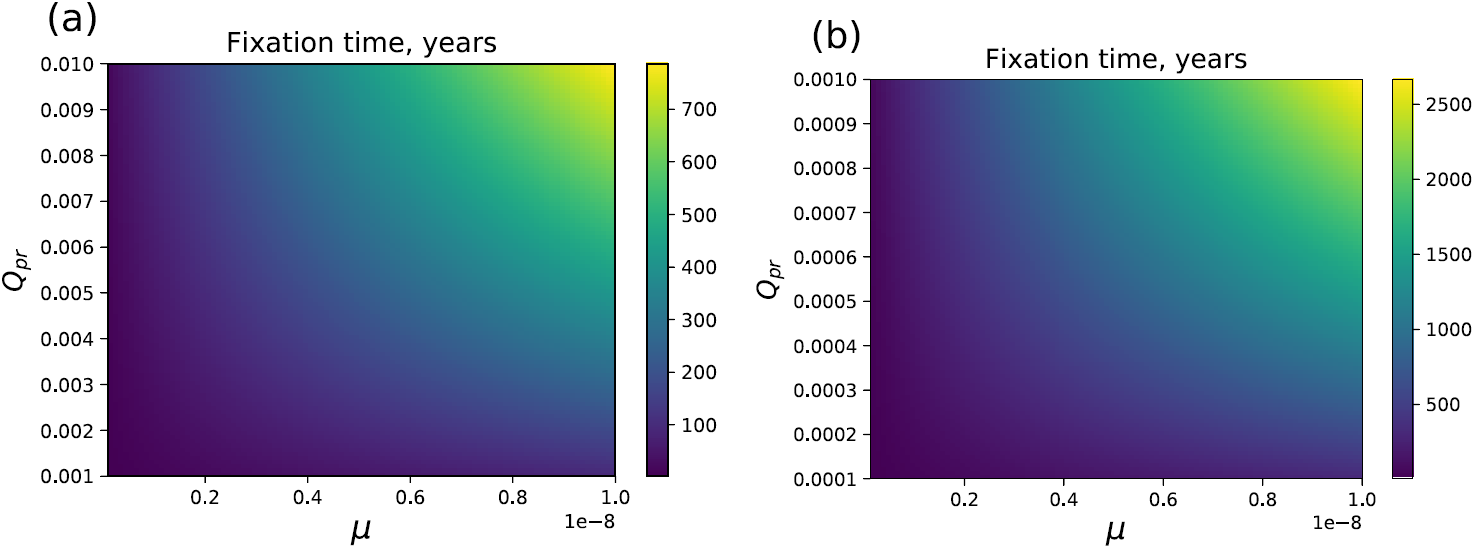
Extinction time *T*_1_ over *μ* − *Q*_*pr*_ parameter space for (a) Colorectal adenocarcinoma (b) Small intestine adenocarcinoma.

